# Structure and substrate specificity determinants of the taurine biosynthetic enzyme cysteine sulphinic acid decarboxylase

**DOI:** 10.1101/2020.09.25.308866

**Authors:** Elaheh Mahootchi, Arne Raasakka, Weisha Luan, Gopinath Muruganandam, Remy Loris, Jan Haavik, Petri Kursula

**Affiliations:** Department of Biomedicine, University of Bergen, Bergen, Norway; Faculty of Biochemistry and Molecular Medicine, University of Oulu, Oulu, Finland; VIB-VUB Center for Structural Biology, Vlaams Instituut voor Biotechnologie, Brussels, Belgium; Structural Biology Brussels, Department of Bioengineering Sciences, Vrije Universiteit Brussel, Brussels, Belgium; Bergen Center of Brain Plasticity, Division of Psychiatry, Haukeland University Hospital, Bergen, Norway; Biocenter Oulu, University of Oulu, Oulu, Finland

## Abstract

Pyridoxal 5′-phosphate (PLP) is an important cofactor for amino acid decarboxylases with many biological functions, including the synthesis of signalling molecules, such as serotonin, dopamine, histamine, *γ*-aminobutyric acid, and taurine. Taurine is an abundant amino acid with multiple physiological functions, including osmoregulation, pH regulation, antioxidative protection, and neuromodulation. In mammalian tissues, taurine is mainly produced by decarboxylation of cysteine sulphinic acid to hypotaurine, catalysed by the PLP-dependent cysteine sulphinic acid decarboxylase (CSAD), followed by non-enzymatic oxidation of the product to taurine. We determined the crystal structure of mouse CSAD and compared it to other PLP-dependent decarboxylases in order to identify determinants of substrate specificity and catalytic activity. Recognition of the substrate involves distinct side chains forming the substrate-binding cavity. In addition, the backbone conformation of a buried active-site loop appears to be a critical determinant for substrate side chain binding in PLP-dependent decarboxylases. Phe94 was predicted to affect substrate specificity, and its mutation to serine altered both the catalytic properties of CSAD and its stability. Using small-angle X-ray scattering, we further showed that similarly to its closest homologue, GADL1, CSAD presents open/close motions in solution. The structure of apo-CSAD indicates that the active site gets more ordered upon internal aldimine formation. Taken together, the results highlight details of substrate recognition in PLP-dependent decarboxylases and provide starting points for structure-based inhibitor design with the aim of affecting the biosynthesis of taurine and other abundant amino acid metabolites.

## Introduction

Pyridoxal 5′-phosphate (PLP), the active form of vitamin B6, is a ubiquitous cofactor essential for a number of enzymes. PLP-dependent enzymes, which mainly use PLP as a covalently bound coenzyme, account for 4% of total cellular enzymatic activity (1, 2). Mammalian genomes encode several PLP-dependent enzymes, which catalyze a variety of biochemical reactions using different substrates (1, 3). Many PLP-dependent enzymes use amino acids as substrates and play central roles in cellular metabolism.

PLP-dependent enzymes are established drug targets in cancer and neurological diseases. Inhibitors of *γ*-aminobutyric acid (GABA) aminotransferase are of therapeutic interest in central nervous system disorders, being used in the treatment of epilepsy (4), and inhibitors of L-DOPA decarboxylase (DDC) are used in treating Parkinson’s disease (5-7). Furthermore, recessive mutations in PLP-dependent enzymes are associated with severe neurological syndromes (8-10). PLP-dependent enzymes can be autoantigens and targets of the immune system in autoimmune disorders, *e.g.* Glu decarboxylase 65 (GAD65; GAD2) in type 1 diabetes (11), GAD67 (GAD1) and GAD65 in several neurological disorders (12), and cysteine sulphinic acid decarboxylase (CSAD) in autoimmune polyendocrine syndrome 1 (13).

The sulphur-containing amino acid taurine is the most abundant free amino acid in mammals (14-16). Taurine is implicated in numerous physiological functions and attracts increasing attention as a biomarker for different disease states (14-16). It has a regulatory role in the maintenance of osmotic pressure and structural integrity of biological membranes (17, 18). In the nervous system, taurine may serve as a growth factor (19, 20) or a neurotransmitter/neuromodulator (21, 22). In many species, taurine deficiency can be lethal or associated with severe disease (17), and in humans, altered levels of taurine have been reported in *e.g.* attention deficit hyperactivity disorder and autism (23). In addition, taurine levels significantly decrease after electroconvulsive treatment in depressed patients, and this decrease strongly correlated with clinical improvements (15, 16). Plasma taurine levels were reduced by 83% in CSAD-deficient mice, and most offspring from 2^nd^ generation *Csad*^*-/-*^ mice died shortly after birth, unless given taurine supplementation (24), indicating a crucial physiological role for CSAD in taurine biosynthesis.

In mammalian tissues, taurine is mainly synthesised from cysteine in a three-step pathway, involving oxidation by cysteine dioxidase (E.C. 1.13.11.20), decarboxylation of cysteine sulphinic acid (CSA) by CSAD (E.C. 4.1.1.29), and oxidation of hypotaurine to taurine. CSA can additionally be decarboxylated by the related enzyme glutamic acid decarboxylase-like protein 1 (GADL1) (25), and taurine can be formed from cysteamine by cysteamine dioxygenase (E.C. 1.13.11.19) (26).

Both CSAD and GADL1 are PLP-dependent decarboxylases (PLP-DC). GADL1, being the closest homologue, displays similar activity as CSAD (25). GADL1 plays a role in the decarboxylation of Asp to β-alanine and, thus, functions in the biosynthesis of the abundant dipeptides anserine and carnosine (27). CSAD and GADL1 have distinct expression patterns in mouse and human brain (25). In the brain, CSAD has been detected in neurons and astrocytes in the cerebellum and hippocampus (28, 29). GADL1 is expressed in muscle, kidney, olfactory bulb, and isolated neurons (25, 30). CSA is the preferred substrate for both CSAD and GADL1, although both are able to decarboxylate cysteic acid (CA) and Asp (25).

In an attempt to understand the molecular mechanisms underlying the catalysis and regulation of GADL1 and CSAD, and to enable future development of optimised inhibitors, we previously solved the crystal structure of mouse GADL1 (31). Here, we determined the crystal structure of mouse CSAD (*Mm*CSAD) in the presence and absence of PLP. Comparisons of the active sites of PLP-DCs highlight substrate recognition determinants within the structurally conserved enzyme family. The structure of CSAD helps to understand the details in the biosynthesis of taurine, one of the most abundant amino acids and dietary supplements.

## Materials and methods

### Expression vector

Multiple mRNA transcripts of CSAD have been described, with alternative initiation codons and splicing events [14]. *Mm*CSAD cDNA corresponding to the 493-amino-acid (55 kDa; UniProt entry Q9DBE0) isoform was subcloned into the expression vector pTH27 (32), which codes for an N-terminal His_6_ tag. In order to examine structural determinants of substrate specificity of *Mm*CSAD, the variant F94S was generated using the QuikChange kit (Agilent). The sequences of expression clones were verified by DNA sequencing.

### Expression and purification of MmCSAD

Wild-type (WT) and F94S His_6_-CSAD were expressed in *Escherichia coli* BL21 CodonPlus (DE3)-RIPL cells (Stratagene) at +15 °C using 0.5 mM IPTG induction. Pyridoxine hydrochloride, a precursor of PLP biosynthesis, was added to the culture at 2 mM to improve protein solubility. Cell pellets were lysed by sonication in a buffer consisting of 50 mM sodium phosphate buffer pH 7.4, 500 mM NaCl, 20 mM imidazole, 0.2 mg/ml lysozyme, 1 mM MgCl_2_, and cOmplete EDTA-free protease inhibitors (Roche). Phenylmethylsulfonyl fluoride was added to 1 mM immediately following sonication, and the unclarified lysate was applied onto an IMAC HiTrap TALON crude column (GE Healthcare). The column was washed first with 50 mM sodium phosphate pH 7.4, 500 mM NaCl, and then with the same buffer containing 20 mM imidazole. Elution was done with 100 mM imidazole in the same buffer. Size-exclusion chromatography (SEC) was performed using a Superdex HR 200 column (GE Healthcare) equilibrated with 20 mM HEPES, 200 mM NaCl (pH 7.4). After elution, CSAD dimer fractions were combined and concentrated.

### Crystallisation, data collection, structure solution, and refinement

*Mm*CSAD crystals were obtained at +20 °C using sitting-drop vapour diffusion. Holo-CSAD crystals were grown in drops containing 200 nl of protein stock (10.3 mg/ml) and 100 nl of reservoir solution (0.15 M KBr, 30% PEG2000MME), and apo-CSAD crystals grew in drops of 100 nl protein and 200 nl reservoir (200 mM Na_2_SO_4_, 100 mM Bis-tris propane, pH 6.5, 20% PEG3350). Crystals were briefly soaked in a cryoprotectant solution containing 80% reservoir solution and 20% glycerol, and flash-cooled in liquid N_2_ prior to data collection.

X-ray diffraction data were collected at 100 K on the automated MASSIF-1 synchrotron beamline at ESRF (Grenoble, France) (33-35). The data were processed and scaled using XDS (36). The structure was solved by molecular replacement in PHASER (37), using the human CSAD crystal structure (PDB entry 2JIS) as template. Refinement was done in phenix.refine (38) and model rebuilding with coot (39). The structures were validated using MolProbity (40). The refined coordinates and structure factors were deposited at the PDB with the entry codes 6ZEK (holo-CSAD) and 7A0A (apo-CSAD).

### Small-angle X-ray scattering

Small-angle X-ray scattering (SAXS) data for *Mm*CSAD were collected on the SWING beamline at the SOLEIL synchrotron (Gif-sur-γvette, France). Scattering was measured in batch mode at three different protein concentrations (1-2.5 mg/ml), from a freshly purified monodisperse dimeric sample. The recorded frames were checked for radiation damage, and data from all concentrations were analyzed to exclude intermolecular events. Data were processed using the beamline software Foxtrot 3.5.2 and analyzed with ATSAS (41). *Ab initio* chain-like models were built with GASBOR (42), normal mode-based conformations were analysed using SREFLEX (43), and the crystal structure was compared to the SAXS data with CRγSOL (44). The SAXS data and models were deposited at SASBDB (45) with the accession code SASDJR7.

### Sequence and structure analysis

Structure superpositions were done with SSM (46). Sequences were aligned with ClustalW (47) and visualised with ESPript (48). Electrostatic surfaces were calculated using APBS and pdb2pqr (49) and visualized in UCSF Chimera (50). PyMOL (Schrödinger) was used for structure visualization and analysis.

### Circular dichroism spectroscopy

Circular dichroism (CD) spectra between 200–260 nm were recorded in triplicate on a Jasco J-810 instrument. Measurements were done at a protein concentration of 0.6 mg/ml in a 1-mm quartz cuvette. The samples were diluted with 10 mM sodium phosphate (pH 7.4), and buffer spectra were subtracted. Thermal denaturation was measured using the CD signal at 222 nm from +25 to +95 °C, at a heating rate of 2 °C/min.

### Differential scanning fluorimetry

A fluorescence-based thermal stability assay (differential scanning fluorimetry, DSF) (51) was performed using a LightCycler^®^480 II instrument (Roche). 20-μl samples were analyzed in 20 mM HEPES, 200 mM NaCl (pH 7.4). CSAD concentration was 0.1 mg/ml, and SγPRO^®^ Orange was used at a 1:1000 (v/v) dilution. The instrument was set to detect emission between 300-570 nm. The heating rate was 2 °C/min from +20 to +99 °C. Five replicates of each sample were measured, using a 384-well plate with an optical film (Roche).

### Quaternary structure analysis

Analytical SEC coupled to multi-angle static light scattering (MALS) was employed to determine the oligomeric states of WT *Mm*CSAD and F94S. SEC was done using the ÄKTA™Purifier FPLC system (GE Healthcare), which was coupled with a RefractoMax 520 module (ERC GmbH, Riemerling, Germany) for measuring refractive index for concentration determination, and a mini-DAWN TREOS light scattering detector (Wyatt Technology). Samples were diluted to 2 mg/ml and centrifuged at 16 000 *g* for 10 min at +4 °C. 200 µg of the protein were applied onto a Superdex 200 Increase 10/300 GL column, pre-equilibrated with 20 mM HEPES, 200 mM NaCl (pH 7.4), at a flow rate of 0.4 ml/min. Astra software (Wyatt) was used for SEC-MALS data analysis.

### Enzymatic activity assays

Catalytic activity of WT *Mm*CSAD and F94S towards the known substrates CSA and Asp was measured at +37 °C, using a reaction mixture of 100 μl containing 6 µM CSAD, 60 mM potassium phosphate (pH 7.4), 5 mM DTT, and 0.5 μM PLP. Adding the amino acid substrate started the reaction. To measure steady-state kinetic properties of the enzymes, 0-50 mM of the substrates were tested. After 60 min, the reaction was stopped by addition of an equal volume of ice-cold ethanol containing 5% acetic acid. For studying GAD activity with Glu as substrate, the reaction mixture of 100 μl contained 60 mM potassium phosphate (pH 7.4), 5 mM DTT, and 2 mM PLP, and the reaction was stopped after 120 min.

The samples were centrifuged at 15700 *g* for 10 min, and the supernatant was transferred onto a microtiter plate and analyzed by HPLC. Samples were diluted with an equal volume of solvent (24% ethanol in 50 mM sodium phosphate, pH 6.0), and 4.2% *o*-phtaldialdehyde (OPA) reagent was added. Samples were injected into a Zorbax Eclipse XDB-C18 column, and the product was determined based on fluorescence detection of the OPA-conjugated amino acid, using excitation at 366 nm and emission at 455 nm. Kinetic parameters were determined by nonlinear regression using the Michaelis–Menten equation in GraphPad Prism 8 (GraphPad Software, La Jolla, California, USA).

## Results and discussion

Taurine, the most abundant free intracellular amino acid in humans, has been implicated in a range of different physiological functions, and it is valuable as an industrial product and dietary supplement. Our aim was to better understand biosynthesis of taurine and the substrate specificity of PLP-DCs, by way of structural characterization of CSAD, which catalyses the conversion of CSA into hypotaurine (Figure 1A).

**Figure 1.**
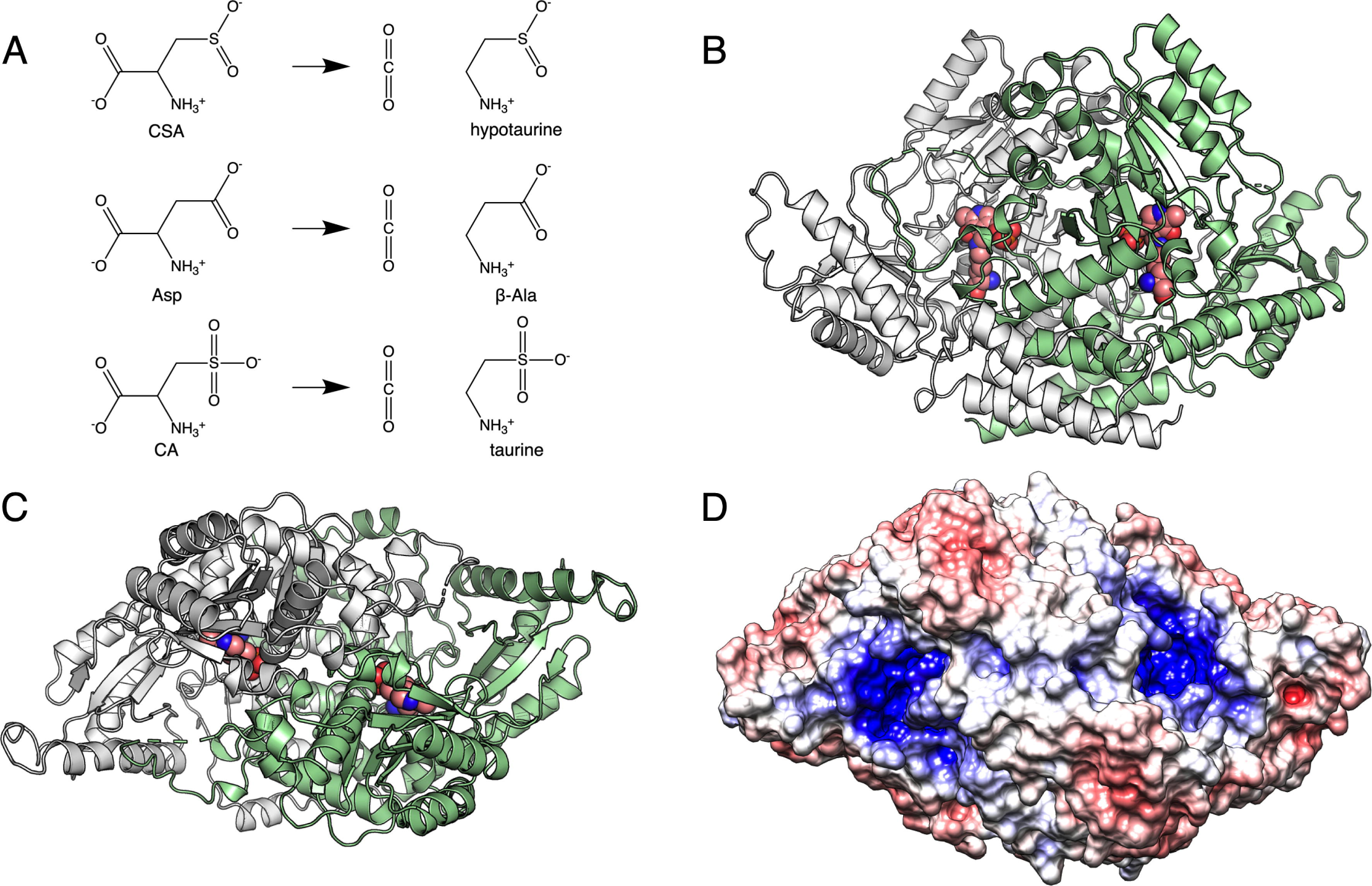
Overall structure of mouse CSAD. A. Reactions catalyzed by CSAD. CSA is the preferred substrate. B. Side view of the dimer, the active site is indicated by the internal aldimine between PLP and Lys305 (spheres). C. Top view of the CSAD dimer. D. Electrostatic surface shows positive potential (blue) in the active-site cavity (top view).

### The crystal structure of MmCSAD

A structure of human CSAD has been available at the PDB (entry 2JIS). However, no detailed information, comparative studies with related enzymes, or mechanistic investigations of CSAD are available. We solved the crystal structure of mouse CSAD, in order to provide a tool for further studies on CSAD catalysis and facilitate the design of small molecules that could be used to modify taurine-related metabolic pathways.

The crystal structure of *Mm*CSAD was solved at 2.1-Å resolution using synchrotron radiation (Table 1, Figure 1B,C). The active site is fully occupied with PLP covalently attached to Lys305 as an internal aldimine, indicating that the structure corresponds to the catalytically competent form of *Mm*CSAD.

**Table 1.** Crystallographic data collection and refinement.

| **Structure** | **CSAD holo** | **CSAD apo** |
| --- | --- | --- |
| Space group | P2<sub>1</sub> | P2<sub>1</sub> |
| Unit cell dimensions | a=72.9, b=113.3, c =113.4 Å<br>$\alpha=\gamma=90^\circ$, $\beta=95.8^\circ$ | a=73.1, b=114.9, c =113.8 Å<br>$\alpha=\gamma=90^\circ$, $\beta=95.8^\circ$ |
| **Data processing** |  |  |
| Resolution range (Å) | 50-2.10 (2.15-2.10) | 50-2.80 (2.87-2.80) |
| Completeness (%) | 90.7 (88.5) | 98.6 (98.6) |
| R<sub>sym</sub> (%) | 12.5 (116.5) | 42.6 (323.1) |
| R<sub>meas</sub> (%) | 15.1 (141.1) | 50.4 (380.9) |
| $\langle I/\sigma(I) \rangle$ | 7.7 (0.9) | 3.4 (0.4) |
| CC<sub>1/2</sub> (%) | 99.3 (38.9) | 94.9 (18.4) |
| Redundancy | 2.7 (2.7) | 3.4 (3.5) |
| Wilson B factor (Å<sup>2</sup>) | 39.2 | 55.1 |
| **Structure refinement** |  |  |
| R<sub>cryst</sub>/R<sub>free</sub> (%) | 17.9/23.5 | 24.3/29.2 |
| RMSD bond length (Å) | 0.007 | 0.003 |
| RMSD bond angle (°) | 0.9 | 0.6 |
| Ramachandran favoured/outliers (%) | 96.3/0.16 | 95.6/0.16 |
| MolProbity score/percentile | 1.61/96<sup>th</sup> | 1.49/100<sup>th</sup> |
| PDB entry | 6ZEK | 7A0A |

The electrostatic potential surface of CSAD (Figure 1D) shows a high positive charge in the active-site cavity, and it is conceivable that this property is important in attracting negatively charged substrates. CSAD and GADL1 prefer amino acids with short acidic side chains as substrates, *i.e.* Asp, CA, and CSA, of which CSA is most favoured for both enzymes (25). On the other hand, Glu, homocysteic acid, and homocysteine sulphinic acid have been reported not to be substrates of CSAD (25, 52).

*Mm*CSAD presents the conserved fold of PLP-DCs, with the closest structural homologues in the PDB being human CSAD and mouse GADL1 (Table 2). A sequence alignment of selected homologues with known structure is shown in Figure 2. High structural similarity was expected, since the chemical reaction catalyzed by PLP-DCs is essentially identical, and the substrates differ from each other only by their respective side chain moieties. It is possible, however, that substrate side chain recognition in different PLP-DCs may cause small changes in the positioning of the reactive groups, thereby leading to different kinetic properties.

**Table 2.**
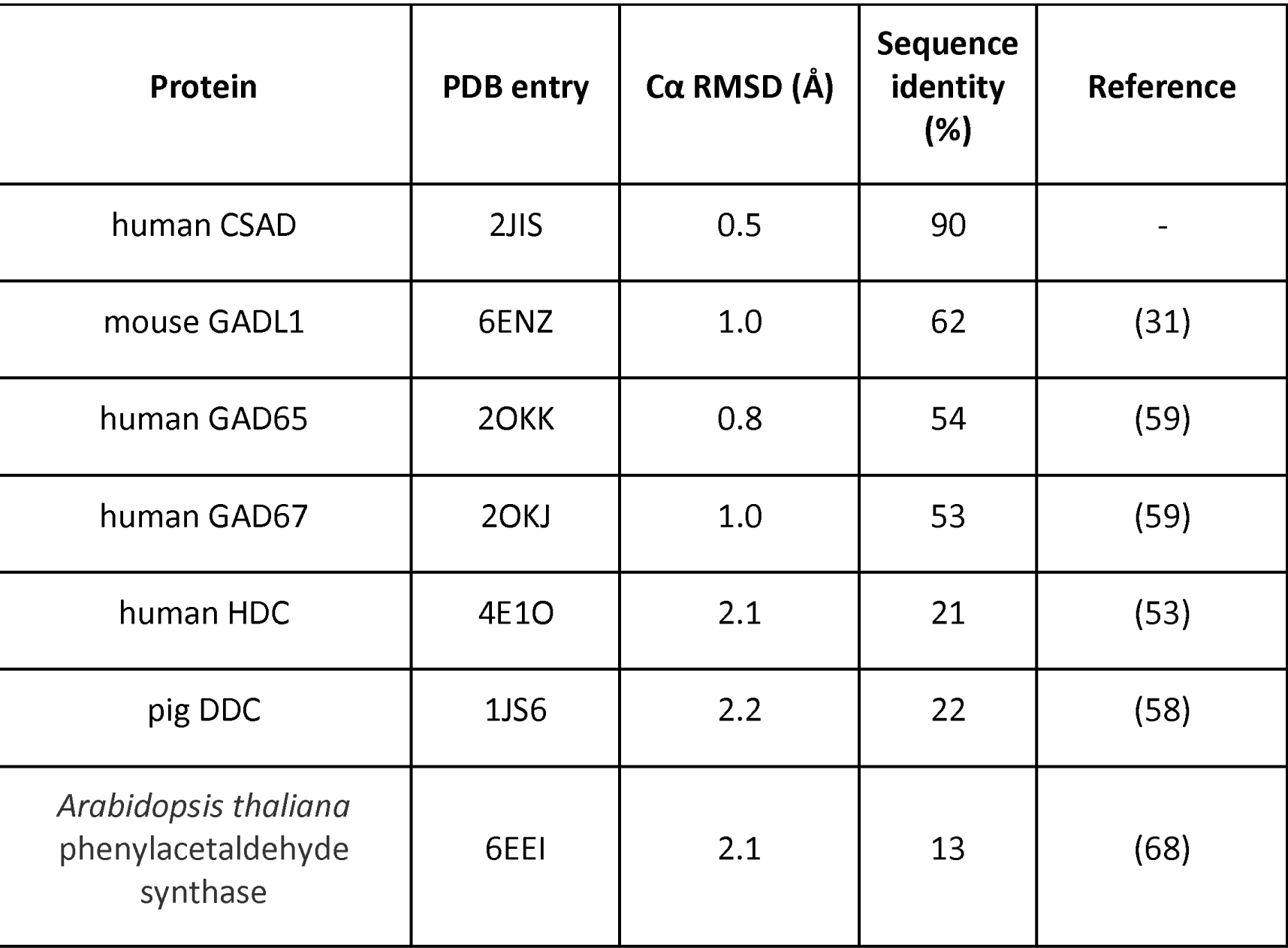
Comparison of the mouse CSAD structure to selected PLP-DCs with known structure. Note how decrease in sequence identity has only minor effects on the similarity of the 3-dimensional fold.

**Figure 2.**
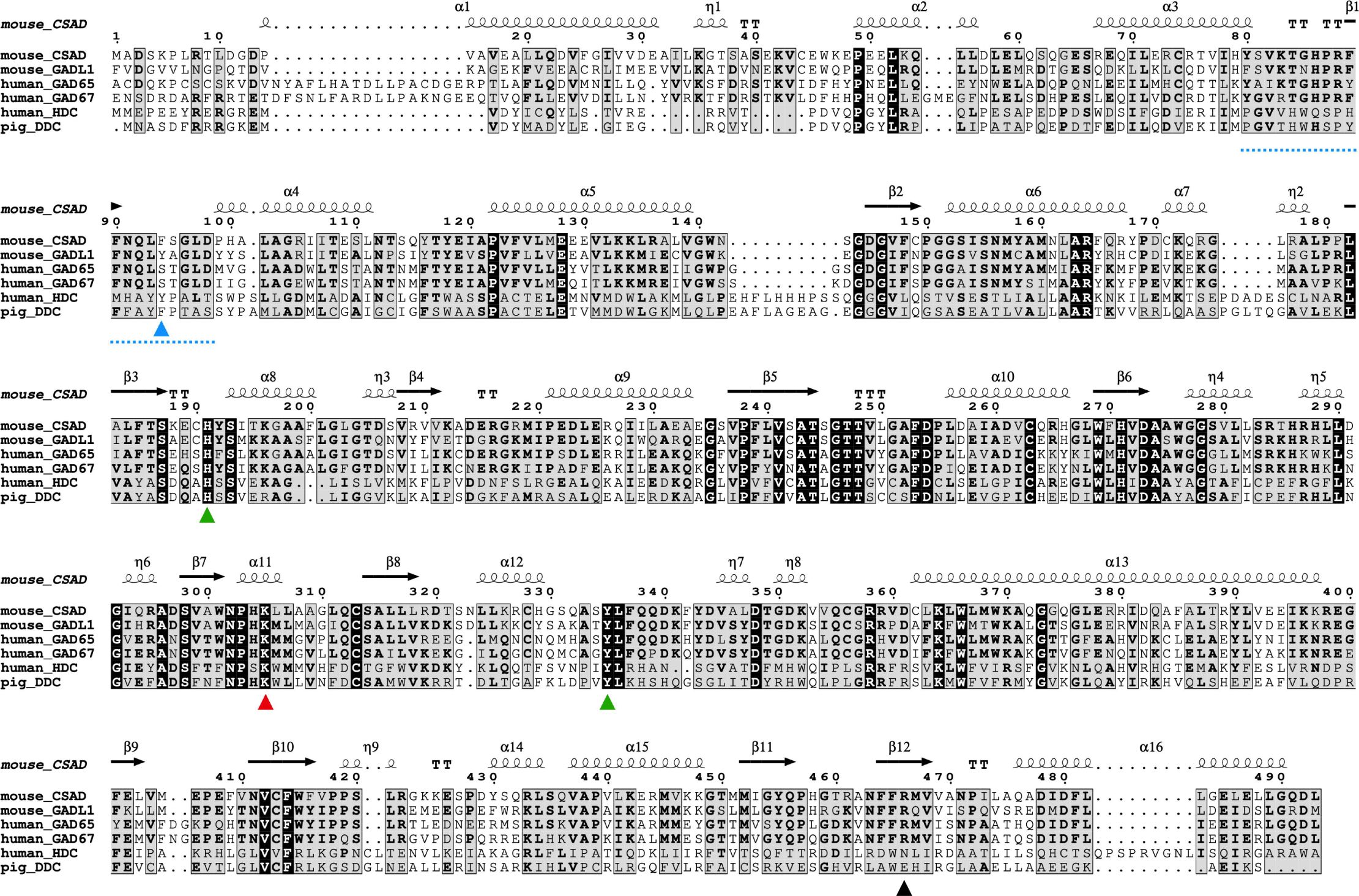
Sequence alignment of PLP-DCs with known structure. Key elements discussed in the text are highlighted: Phe94 (blue), α3-α4 loop (blue dash), His191 (green), Lys305 (red), Tyr335 (green), Arg466 (black).

In addition to the catalytically crucial residue Lys305, which in all PLP-DCs forms the internal aldimine with PLP, other residues in the active site are relevant for substrate binding and catalysis (Figure 3A). The residues interacting with PLP are highly conserved and include His191, which is stacked above the PLP aromatic ring. His191 is fully conserved in the PLP-DC family, and in addition to fixing PLP in a reactive conformation, its roles have been suggested to be central in coordinating the carboxyl group that will be released as CO_2_ (53), as well as in protonating the quinonoid intermediate resulting from decarboxylation (3, 54).

**Figure 3.**
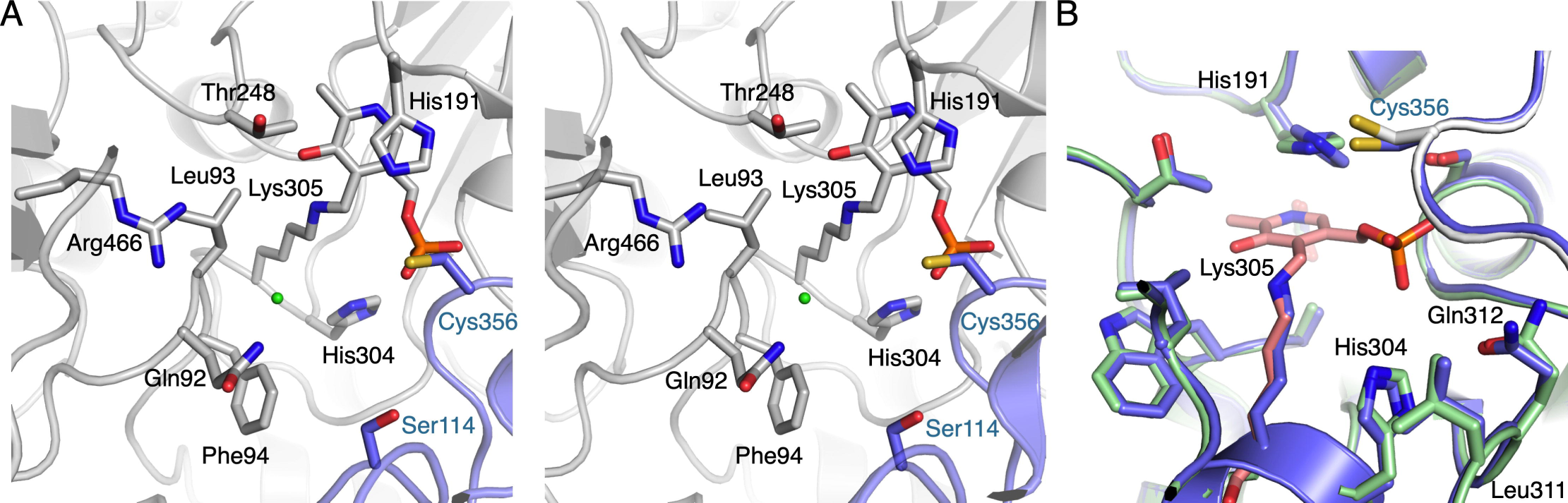
The active site of CSAD. A. Stereo view of the CSAD active site. The bound chloride ion is shown as a green sphere. Elements coming into the active site from the opposing monomer are in blue. B. Comparison of the holo (green/pink) and apo (blue) CSAD active sites.

An additional crystal structure was solved and turned out to be the apo form of CSAD, with no electron density for PLP in the active site (Supplementary Figure 1). Hence, this structure represents an inactive form of CSAD. The main difference between the apo and holo forms is the rotation of the His191 side chain to a conformation not compatible with the presence of PLP (Figure 3B). Overall, the active site of CSAD is less ordered in the absence of PLP; thus, PLP binding will stabilize a catalytically competent conformation of the cavity. No larger-scale conformational changes were observed between the apo and holo enzymes, which is different from DDC, in which the structure of the apo form presented an open conformation, suggested to be linked to the mechanism of cofactor loading (55). Interestingly, both CSAD structures were obtained from the same protein batch, indicating that PLP has been lost during crystallisation of the apo form. One possibility is hydrolysis of the internal aldimine at the slightly acidic pH (6.5) of the conditions giving the apo CSAD crystals, eventually leading to loss of PLP from the active site.

### Structure of CSAD in solution

In addition to the crystal structure, we studied CSAD conformation in solution using synchrotron SAXS (Figure 4), in order to detect possible flexibility, as previously observed for GADL1 (31). Indeed, dimeric CSAD behaves much like GADL1 in solution, providing evidence for motions between open and closed states of the dimer. Such motions could be related to the catalytic cycle. They could be linked to the conformation of the flexible catalytic loop (residues 330-340), which is not fully visible in the CSAD crystal structure. Like in GADL1 (31), normal mode analysis identified an open conformation of the CSAD homodimer, which fits the SAXS data well (Figure 4), but is different from that observed crystallographically for apo-DDC (55). Whether open/close motions are unique to each class of PLP-DCs, or if all family members are equally dynamic, remains a subject for future research.

**Figure 4.**
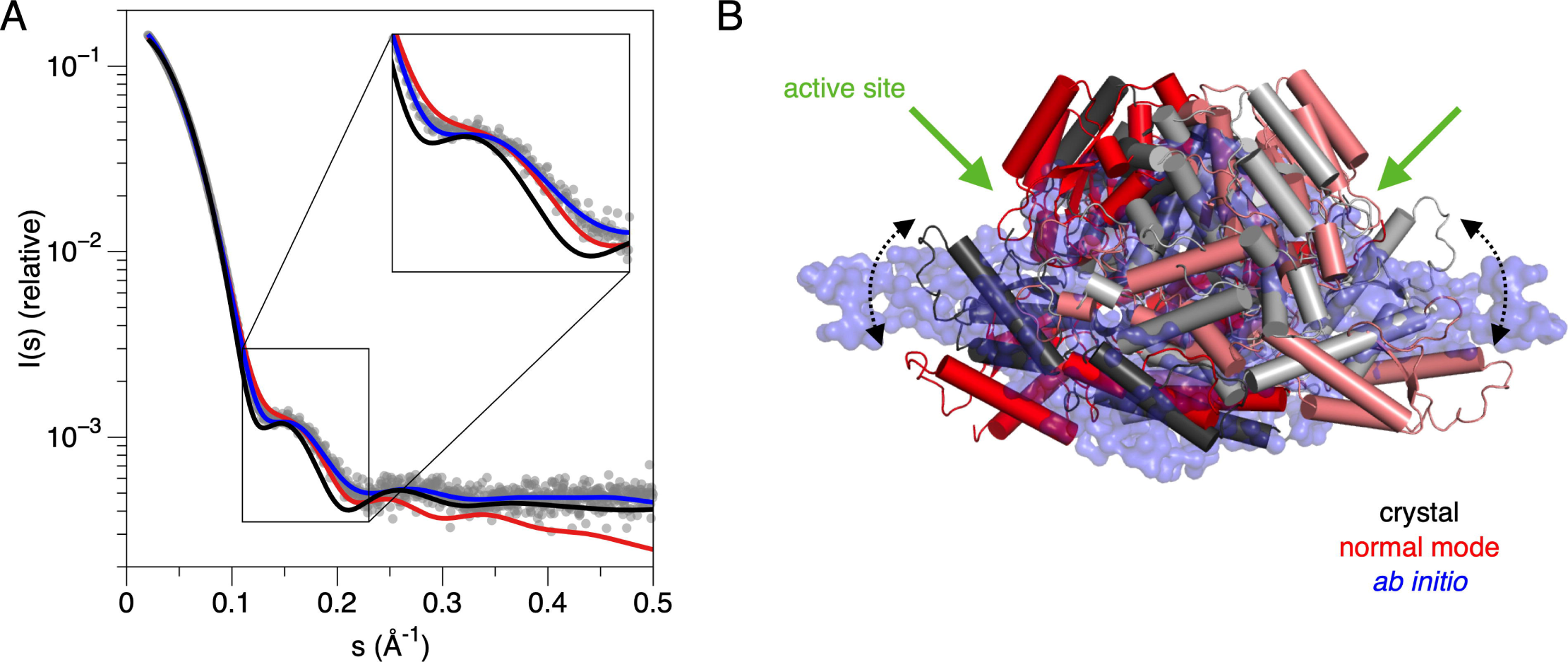
Structure of CSAD in solution. A. SAXS data (grey dots) overlaid with fits from the crystal structure (black), the *ab initio* model (red), and the normal mode-based conformation (blue). B. Comparison of the crystal structure and models. The open/close motions and the active site location are indicated.

### Importance of Phe94 for substrate specificity

When comparing the structures of CSAD and GAD, it can be concluded that Phe94 in CSAD may be important for substrate specificity. Apparently, Phe94 blocks the binding site for larger substrates; hence, Glu should not productively bind (Figure 5A). The active site of GAD has a Ser residue at the corresponding position. In addition, GADL1 has a Tyr residue at this position, linked to a slightly different substrate preference (25). We previously identified this position as a key difference in the substrate recognition pocket in the otherwise highly homologous acidic amino acid decarboxylases (27). Thus, we mutated Phe94 to Ser in CSAD to evaluate the effects on enzymatic properties.

**Figure 5.**
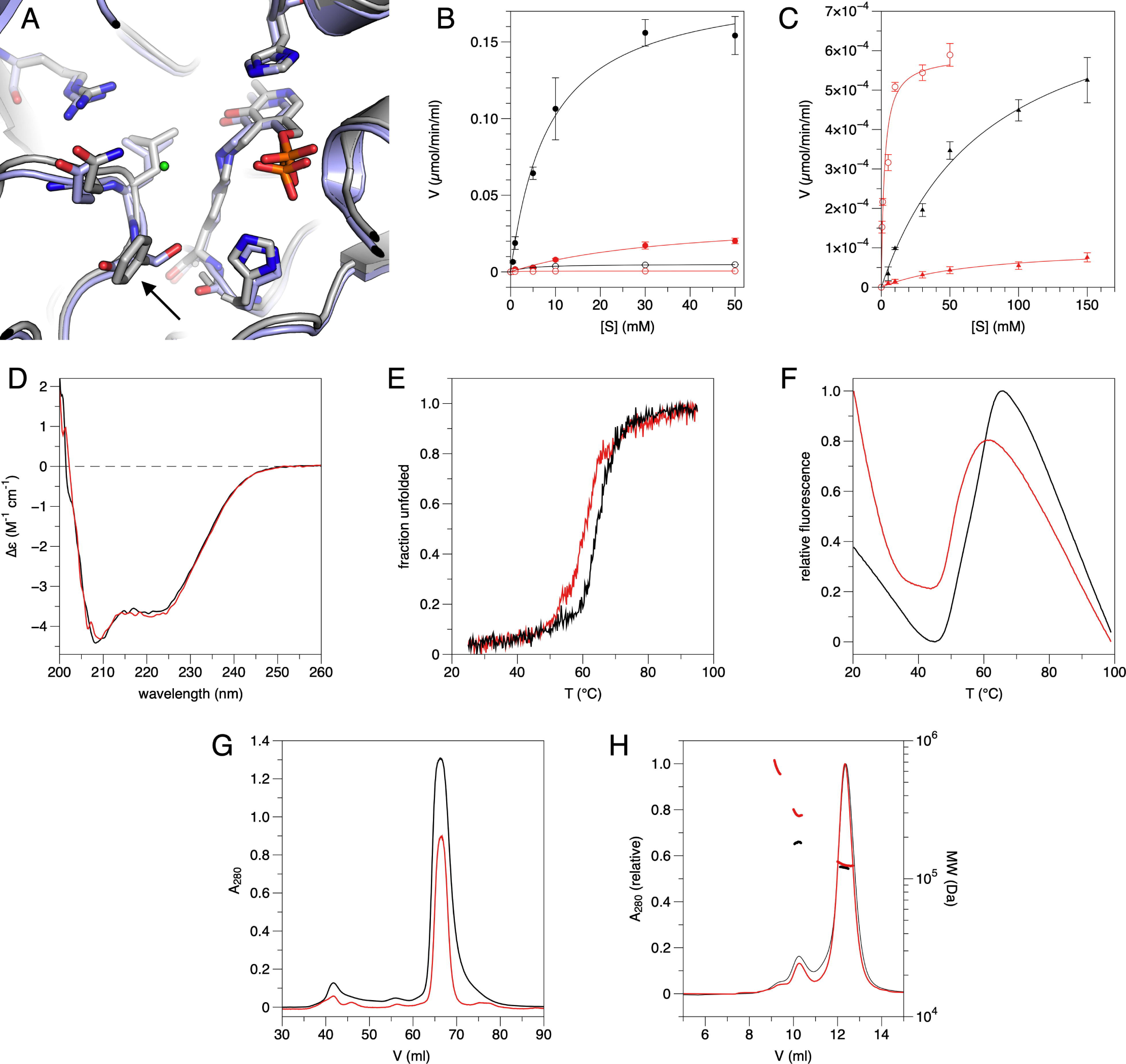
Role of Phe94 in CSAD activity and stability. A. Superposition of CSAD (grey) and GAD67 (light blue) active sites. The arrow indicates the position of CSAD Phe94. The chloride ion in CSAD is shown in green. B. Activity assay with CSA (filled symbols) and Asp (open symbols). WT, black; mutant, red. C. Activity assay with Glu (filled triangles). WT, black; mutant, red. The activity level of the mutant with Asp is shown for reference (red open circles). D. CD spectra for WT (black) and mutant (red) CSAD. E. CD melting curves. F. DSF melting curves. G. SEC during protein purification; the pure dimer peak at ∼67 ml was picked for further experiments. H. SEC-MALS after freezing and thawing of dimeric CSAD indicates presence of some higher-order oligomers in both WT and F94S.

Activity assays of WT and F94S *Mm*CSAD towards CSA and Asp were carried out using HPLC (Table 3, Figure 5B). While the F94S mutation affected both K_m_ and k_cat_ of CSAD towards CSA and Asp, the mutant enzyme remained active. Specifically, the effects on K_m_ and k_cat_ revealed that F94S has a turnover number 5-10 times lower than WT CSAD for both CSA and Asp, indicating an overall effect on catalysis. In the mutant, K_m_ increases by an order of magnitude for CSA, but not for Asp. Hence, Phe94 is specifically important for the binding of CSA, the preferred substrate of CSAD. k_cat_/K_m_ values further support these observations, showing that the most effective combination by far is WT CSAD with CSA as substrate.

**Table 3.** Enzymatic properties of CSAD towards acidic amino acids.

| Substrate | WT CSAD |  |  | F94S |  |  |
| --- | --- | --- | --- | --- | --- | --- |
| | $k_{\text{cat}}$ ($\text{s}^{-1}$) | $K_m$ (mM) | $k_{\text{cat}}/K_m$ ($\text{M}^{-1}\text{s}^{-1}$) | $k_{\text{cat}}$ ($\text{s}^{-1}$) | $K_m$ (mM) | $k_{\text{cat}}/K_m$ ($\text{M}^{-1}\text{s}^{-1}$) |
| CSA | $0.53 \pm 0.02$ | $8.6 \pm 1.2$ | 61.6 | $0.10 \pm 0.01$ | $36.8 \pm 8.5$ | 2.7 |
| Asp | $0.014 \pm 0.004$ | $3.8 \pm 0.5$ | 3.7 | $0.0016 \pm 0.00006$ | $2.3 \pm 0.4$ | 0.7 |
| Glu | $0.0022 \pm 0.0002$ | $77.3 \pm 12.2$ | 0.03 | $0.00029 \pm 0.00004$ | $72.6 \pm 20.9$ | 0.004 |

A test for GAD activity was carried out using Glu as substrate, as the mutation F94S mimics the GAD substrate binding site (Figure 5C). While a trace level of activity is observed for WT CSAD, this activity is even weaker for F94S. The K_m_ values for both variants are an order of magnitude higher than for CSA and Asp. Hence, altering the substrate specificity towards Glu requires more than altering the obvious Phe94 to the corresponding Ser residue of GAD. Additional factors may include protein dynamics, effects of the catalytic loop – not visible in any structure of CSAD or GADL1 – as well as minor conformational changes in the active site caused by the mutation.

### Folding and stability of WT MmCSAD and F94S

The folding and thermal stability of WT *Mm*CSAD and F94S were examined using DSF and CD spectroscopy. WT *Mm*CSAD and F94S show CD spectra with two characteristic minima at 208 and 220 nm. Both spectra are essentially identical, indicating correct folding of the mutant (Figure 5D). Both CD spectroscopy and DSF indicate decreased thermal stability for the F94S mutant (Table 4). From the CD melting curves it follows that although WT *Mm*CSAD and F94S have similar patterns of melting, F94S has a melting temperature ∼5 °C lower than the WT protein (Figure 5E). While the DSF melting curves indicate a similar decrease in stability for F94S (Figure 5F), the mutant appears to open up already at low temperatures. The differences in T_m_ between the methods, which are commonly seen, could be because of differences in the measurement method of CD and DSF. While CD measures the unfolding of secondary structures based on peptide backbone conformation, DSF follows access of a small-molecule dye to the protein hydrophobic core.

**Table 4.** Thermal stability of WT and F94S *Mm*CSAD.

| Protein | DSF $T_m$ ($^{\circ}\text{C}$) | CD $T_m$ ($^{\circ}\text{C}$) |
| --- | --- | --- |
| *Mm*CSAD | $56.0 \pm 0.2$ | $64.6 \pm 0.1$ |
| F94S | $51.1 \pm 0.2$ | $60.5 \pm 0.2$ |

The effect of the F94S mutation on the oligomeric state and long-term stability of *Mm*CSAD was further studied using SEC-MALS, after freezing and thawing of a pure dimer fraction (Figure 5G,H). Both WT *Mm*CSAD and F94S displayed very similar elution profiles, which mainly correspond to a dimer (110 kDa). Both variants also have some tetramer and higher-order oligomers after freezing and thawing (Figure 5H). Notably, no monomeric form was detected, indicating high-affinity dimerization; as the active site is formed at the dimer interface, this is expected.

Taken together, the F94S mutation does not affect the secondary structure content of CSAD, but it has a clear effect on protein stability. As the mutant displays lower activity than the WT enzyme, at least part of the effect could be from altered protein dynamics. These data indicate an important role for Phe94 in CSAD substrate binding and activity.

### Comparison of the mouse CSAD active site with other decarboxylases

As far as enzymatic activity is concerned, CSAD is known to prefer CSA as substrate, while having weak activity towards CA and Asp (25, 52). The closest homologue, GADL1, is more active towards Asp than CSAD, although it is most active with CSA as substrate. Based on earlier studies, neither of the enzymes accept Glu as substrate, and Glu fails to act as an inhibitor (25), indicating lack of binding to the active site. It is possible that close members of the PLP-DC enzyme family have at least partially overlapping activities. Based on the observation that mice lacking GADL1 have organ specific reductions of β-alanine and taurine levels, we recently suggested that this enzyme might have multiple physiological substrates also *in vivo* (27). On the other hand, although our results indicate Glu is a very weak substrate for CSAD and GADL1, this activity is unlikely to be of any physiological relevance.

Using the high-resolution crystal structures of both CSAD and GADL1, as well as other PLP-DCs (Table 2), one can carry out an inspection of the corresponding active-site geometries underlying substrate specificity. As all PLP-DCs catalyse the same chemical reaction, amino acid decarboxylation, differences in the active site are expected to affect substrate specificity and/or affinity, rather than reaction mechanism *per se*. Hence, catalytically crucial features are expected to be conserved in sequence and 3D structure. For example, His191 is mechanistically important in PLP-DCs, possibly being involved in the protonation of the quinonoid reaction intermediate (3). Comparison of the apo and holo CSAD structures indicates that the conformation of His191 is linked to the presence of the PLP cofactor; it can be envisaged that His191 is important for the catalytically competent orientation of PLP and *vice versa*.

A comparison of CSAD and GADL1 should give indications on the structural properties causing their differential preference towards Asp, although CSA is the preferred substrate for both. Asp and CSA have slightly different geometries, in that the carboxyl group is planar while the sulphinic acid group is not. In addition, for the third known substrate, CA, the molecular size is larger, with 3 oxygen atoms interacting with the recognition pocket. In the predicted binding pocket for the acidic side-chain group, Phe94 in CSAD is replaced by Tyr in GADL1; while the hydroxyl group points away, minor changes in conformation and/or dynamics could explain the differences between CSAD and GADL1. The side chain and backbone of Glu92, Leu93, and Phe94 are likely to define the binding of the side chain in CSAD substrates (Figure 5A).

When comparing CSAD and GAD65/67, it becomes obvious that Phe94 in CSAD is important for substrate specificity. It blocks the binding site for larger substrates, and the Ser residue at this position in GAD is likely to form H-bonds to the Glu side chain carboxyl group (Figure 6A). The binding mode of Glu into GAD can be deduced from the structure of GAD65 in complex with the inhibitor chelidonic acid (Figure 6A) (56). Original studies on this line of inhibitors indicated that the distance between the two carboxyl groups, corresponding to an extended Glu molecule, is important (57). The carboxyl groups of the inhibitor mimic those of Glu, and H-bonds are seen to Ser183/192 and a water molecule coordinated by His395/404 at the bottom of the cavity (numbering for GAD65/67), and well as to the backbone amide groups of the α3-α4 recognition loop containing Ser183/192 (see below). The His residue and the water molecule are also present in CSAD, and CSA is likely to similarly H-bond to this water molecule.

**Figure 6.**
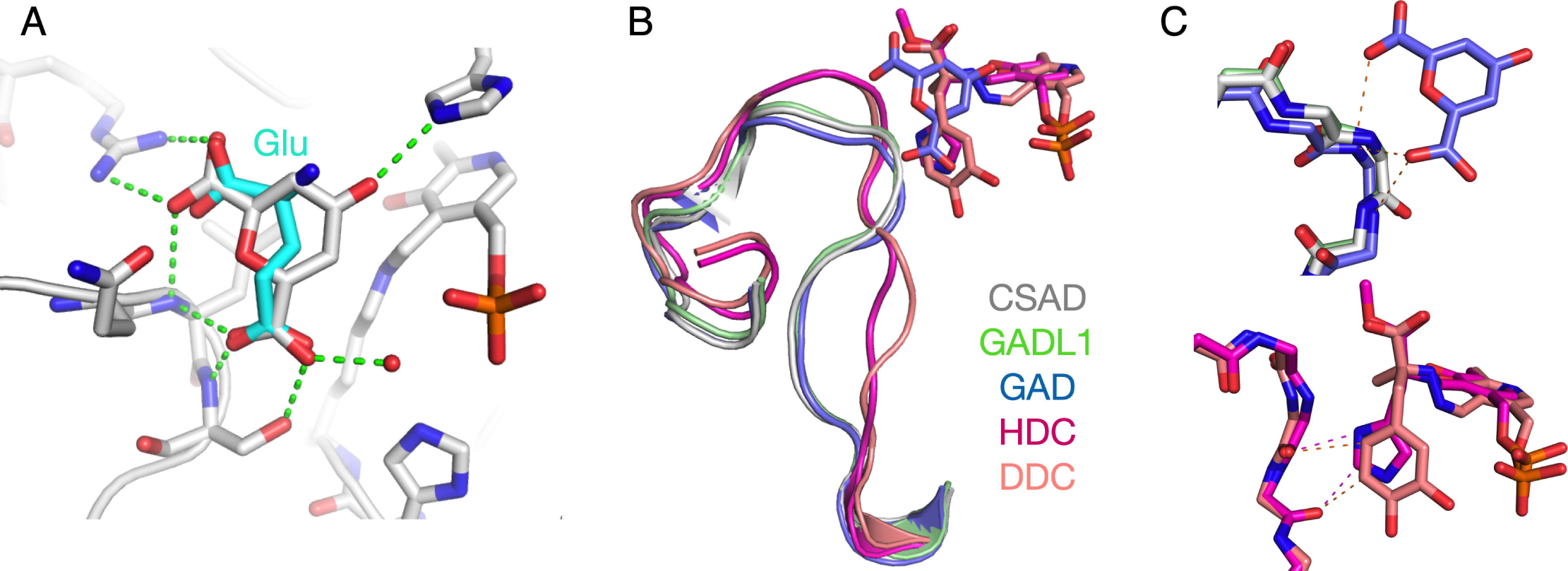
Comparison to other decarboxylases. A. GAD65 in complex with chelidonic acid, with hydrogen bonds shown in green. Based on the structure, a proposed mode of Glu substrate binding in GAD is shown (cyan). B. Superposition of the α3-α4 loop in CSAD (grey), GADL1 (green), GAD65 (blue), HDC (magenta), and DDC (pink) highlights two subfamilies linked to substrate side chain size and properties. C. Top: Backbone conformation in acidic amino acid decarboxylases (with chelidonic acid). Amino groups from the α3-α4 loop form direct hydrogen bonds (dashes) with the acidic substrate. Bottom: Backbone conformation in HDC and DDC; carbonyl groups interact with the substrate *via* hydrogen bonds and/or van der Waals interactions (dashes).

HDC has been crystallized with a non-cleavable methylated analogue of His as a PLP adduct, thereby giving insights into the reaction mechanisms in the family (53). A similar conformation was observed for the inhibitor carbidopa in DDC (58). Phe94 in CSAD prevents the binding of such large substrates into the active site, but it is, in fact, not the only determinant. Comparing the acidic amino acid decarboxylases GAD65, GAD67, CSAD, and GADL1 with those acting on larger, non-acidic substrates, one can observe the α3-α4 loop, forming the active site cavity wall and intimately interacting with the substrate (Figure 6B). The backbone conformation of the loop is different between the two groups of enzymes, and the peptide bonds have opposite orientations, such that the NH groups point towards the substrate-binding cavity in GAD, CSAD, and GADL1 (Figure 6C). In HDC and DDC, the carbonyl groups point into the same pocket, giving a different electrostatic environment and allowing the binding of positively charged and neutral substrates, such as His or L-DOPA, into the active site (Figure 6C).

### Proposed substrate binding mode and catalytic mechanism in CSAD and GADL1

A number of ions were detected in the *Mm*CSAD structure; including an ion modelled as chloride in each active site (Figure 3A,5A), interacting with the backbone amide groups of loop α3-α4 discussed above. This site could be relevant for recognizing the substrate acidic side chain. In the structure of HDC with methylhistidine (53) covalently linked to the PLP cofactor, the binding determinants for the carboxyl and amino groups are well defined, while the inhibitor chelidonic acid bound to GAD (56, 59) provides additional information. Using these structures as templates, one can dock in CSA, taking into account the anion-binding site in the active cavity (Figure 7). We consider the latter to be a likely location for the CSA sulphinic acid group binding - or in the case of Asp, the side-chain carboxyl group.

**Figure 7.**
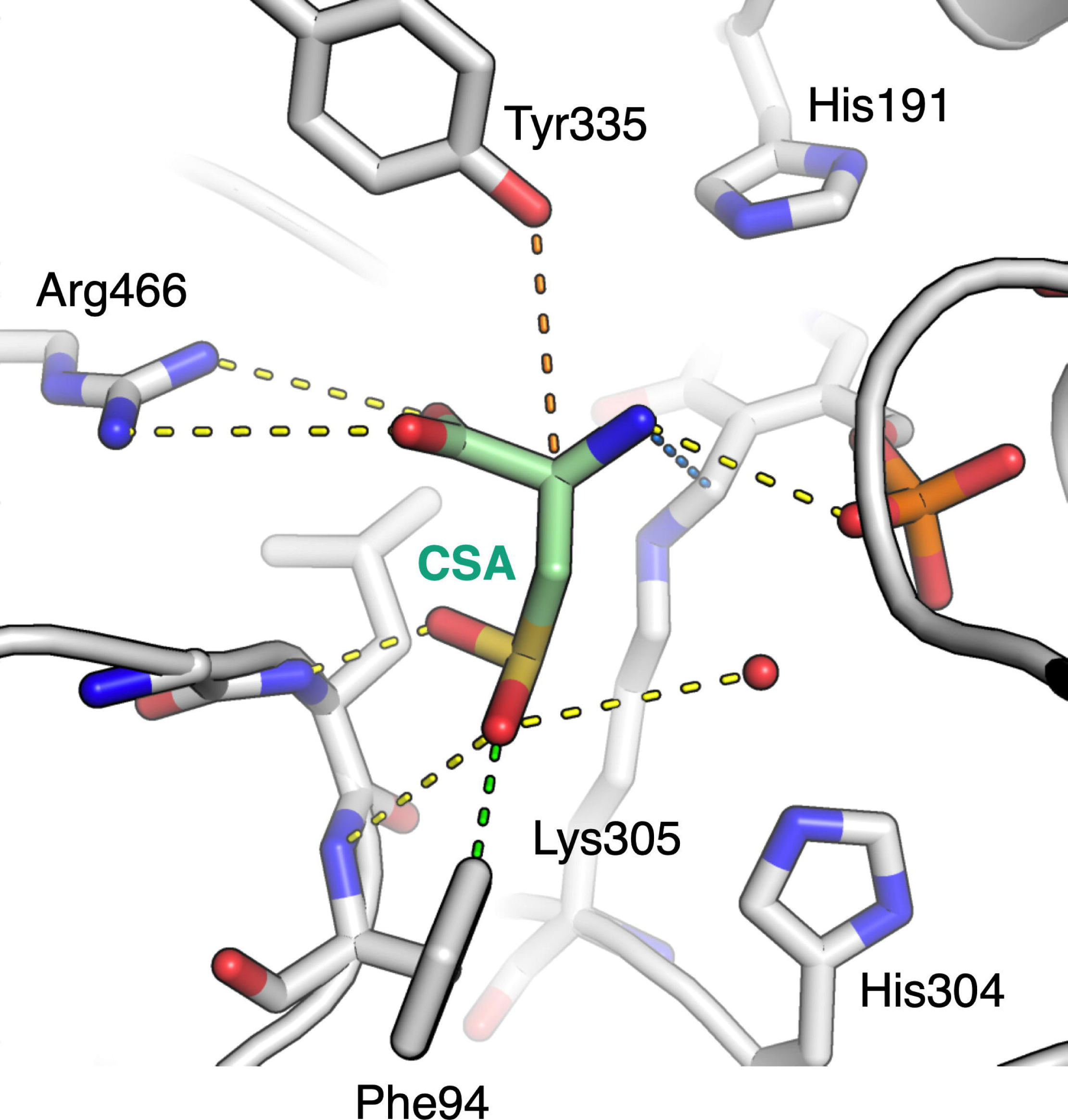
Proposed model for CSA substrate binding to CSAD and depiction of catalytically important residues. Hydrogen bonds to active-site side chains, main-chain groups, the cofactor, and a conserved water molecule are shown in yellow dashes. A van der Waals interaction possibly relevant for CSAD substrate specificity to Phe94 is in green. The blue dash indicates the bond that will be formed upon external aldimine formation between Lys305 and CSA. Protonation of the quinonoid intermediate at the second stage of the reaction is likely catalyzed by Tyr335 from the flexible α12-α13 catalytic loop (here pictured in the position observed in the GAD65 crystal structure), from the other subunit.

The leaving carboxyl group, in a perpendicular position to the plane formed by the PLP ring and the Schiff base moiety, is stabilized by Arg466 (Figure 7). It may be also be H-bonded to His191, a central residue in the PLP-DC mechanism, as seen in the structure of HDC (53). In apo CSAD, His191 is rotated, and it will reach its correct conformation upon PLP complex formation. Thus, the core active site of CSAD appears pre-organised for catalysis, but only in the presence of the cofactor internal aldimine.

A main obstacle in fully understanding the mechanistic details of CSAD and GADL1 catalysis is the flexible α12-α13 loop covering the active site; this loop is not visible in any of the available CSAD or GADL1 crystal structures, but a conserved Tyr residue in it has been suggested to be a key player in catalysis by PLP-DCs, including GAD (59). This Tyr, Tyr335 in CSAD, is conserved in CSAD and GADL1 (Figure 2). Importantly, in both DDC and HDC, the loop is susceptible to proteolysis, but gets protected, when an active-site ligand is bound (58, 60, 61). Hence, the dynamics of the loop are linked to the decarboxylase reaction cycle and occupancy of the active site. After decarboxylation, protonation of the reaction intermediate is carried out by Tyr335, which enables the reaction to proceed towards product release and reconstitution of the internal aldimine in the CSAD active site. The protonation could either occur directly (59) or be mediated through a water molecule coordinated by Tyr335 (62). Intriguingly, PLP-DCs can catalyse a different reaction involving molecular oxygen, if this Tyr residue is mutated to a Phe (63, 64). His191 also appears to be relevant for the protonation step (3, 54, 62), but in light of current data, its likely role is the coordination of Tyr335.

### Insights into inhibitor design

Rare mutations in enzymes responsible for the degradation of β-alanine, as well as its histidine derivatives carnosine and anserine, have been linked to the abnormal accumulation of these compounds in mammalian tissues and severe neurological diseases (65, 66). As the neurological symptoms have been unresponsive to dietary interventions, an alternative treatment strategy could be to inhibit their biosynthesis. The discovery (27) that GADL1 functions in the biosynthesis of β-alanine and carnosine raises the possibility that it could be a target for inhibition therapy. The first generation of inhibitors targeting CSAD and GADL1 had modest affinity but promising selectivity (25). Since then, high-resolution structures for both enzymes have become available, providing a stepping stone for further knowledge-based optimization of such compounds. The differences in activity towards closely related substrates as well as the subtle differences in the active-site structures can be utilised in the development of the next generation of potential inhibitors of acidic amino acid decarboxylases. These aspects are crucial in the development of *in silico* screening approaches.

The prime example of inhibitor design towards PLP-DCs is carbidopa (7), which is a DDC substrate analogue able to form a stable covalent adduct with the PLP cofactor. Carbidopa is in wide use in the treatment of Parkinson’s disease, whereby it inhibits the conversion of L-DOPA to dopamine in peripheral tissues (6), leading to increased L-DOPA half-life and reduced side effects of dopamine. The crystal structure of DDC in complex with carbidopa (58) is therefore of value in designing inhibitors towards other PLP-DCs. For example, for acidic amino acid decarboxylases, a similar approach targeting the external aldimine instead of a simple substrate analogue – as done previously (25, 57, 67) – would be a promising approach. Like carbidopa, non-cleavable analogues of different stages of the acidic amino acid decarboxylase reaction mechanism could be targeted in a systematic manner instead of non-targeted screening or attempts at designing substrate analogues, which inherently will have rather low affinity in these enzymes.

## Concluding remarks

Our work highlights important details on the molecular mechanisms of the biosynthesis of taurine; in addition, substrate recognition determinants across the PLP-DC family have been elucidated. Substrate specificity in the family is clearly affected by both the amino acid side chains lining the catalytic cavity as well as direct backbone-substrate interactions. The latter divide PLP-DCs into two subclasses. These findings are central in understanding mechanistic details of catalysis, but also in research aimed at designing effectors of amino acid decarboxylation linked to the production of important metabolites and signalling molecules, such as taurine, β-alanine, carnosine, GABA, histamine, serotonin, and dopamine.

---

**Supplementary Figure 1. Electron density maps for the holo (top) and apo (bottom) forms of CSAD.** Shown are the final refined 2F_o_-F_c_ maps contoured at 1 σ. Both maps are shown in stereo view, and key residues either changing conformation or becoming more flexible in apo CSAD are labelled.

## Acknowledgements

This work has received funding from the European Union Horizon 2020 research and innovation program under Grant Agreement No. 810384 (CoCA), Stiftelsen Kristian Gerhard Jebsen (SKJ-MED-02), and the Regional Health Authority of Western Norway (No. 25048). We wish to acknowledge access to and excellent support on synchrotron beamlines at SOLEIL and ESRF.

## Notes

### Competing Interest Statement

The authors have declared no competing interest.

