## Supplementary figures and images for "Structure and substrate specificity determinants of the taurine biosynthetic enzyme cysteine sulphinic acid decarboxylase"

### Supplementary Figure 1

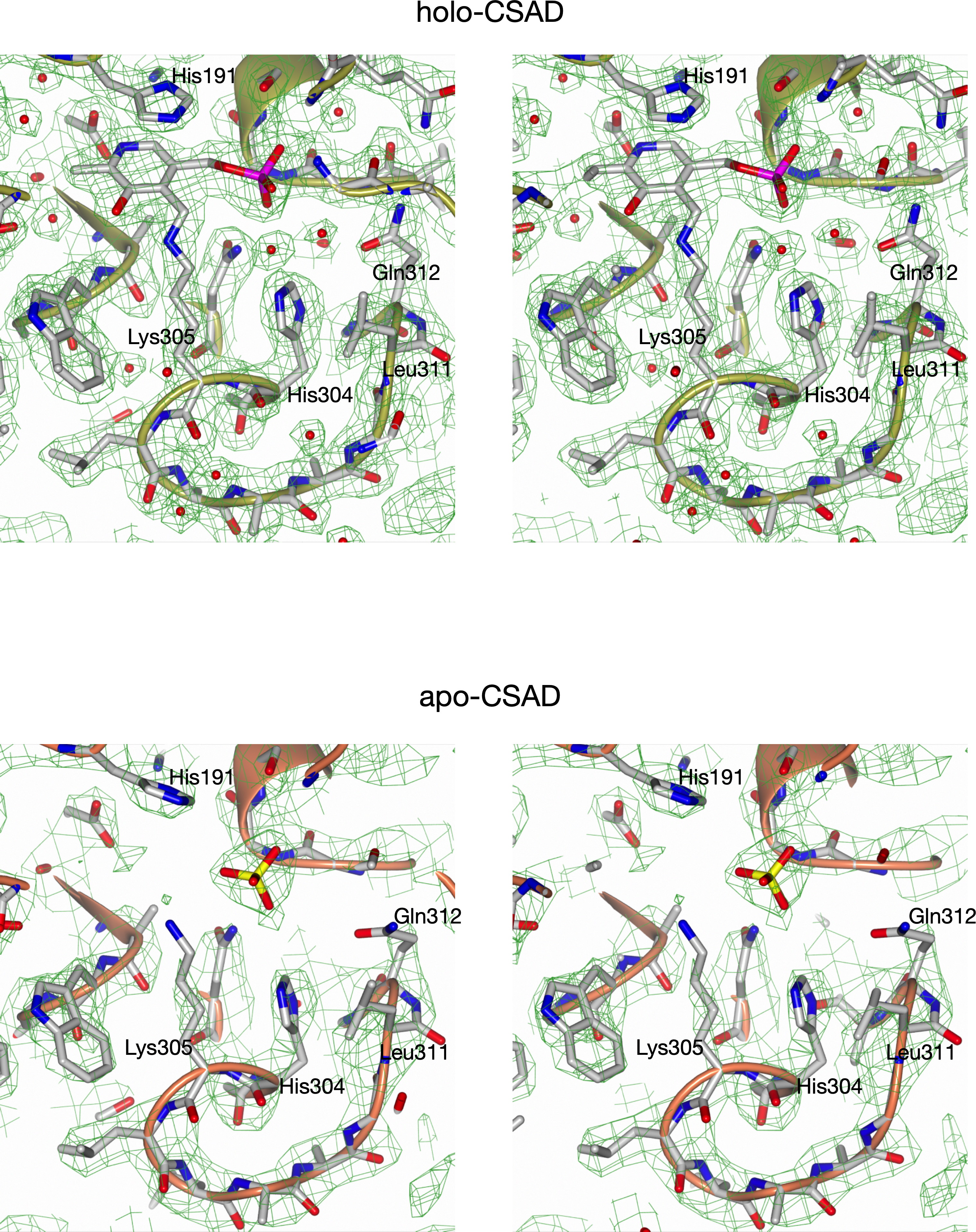
